# Elucidating the Type 2 Diabetes Regulatory Network: Identification of EGFR as a Key Hub and Novel Drug Candidates

**DOI:** 10.1101/2024.07.08.602441

**Authors:** Ricardo Romero Ochoa, Celic Abigail Cohen Rojas

## Abstract

**Introduction:** Type 2 diabetes (T2D) is a complex metabolic disorder with incompletely understood molecular mechanisms. This study aimed to elucidate the T2D regulatory network and identify potential drug targets and candidates.

**Methods:** We performed differential gene expression analysis on multiple T2D datasets, constructed protein-protein interaction networks, and conducted a meta-analysis to identify key hub genes. Functional enrichment analysis was performed on the resulting network. Structure-based virtual screening targeting EGFR, followed by molecular dynamics simulations, was used to identify potential drug candidates.

**Results:** EGFR emerged as a consistently top-ranked hub gene across studies. The regulatory network comprised hub genes, transcription factors, and miRNAs involved in processes such as apoptosis regulation, cellular response to organic substances, and reactive oxygen species metabolism. Virtual screening identified three compounds with favorable ADMET properties and strong binding affinities to EGFR, outperforming control drugs. These compounds demonstrated stable interactions in molecular dynamics simulations.

**Conclusions:** Our integrative analysis provides new insights into the T2D regulatory network, highlighting EGFR as a potential therapeutic target. The identified drug candidates offer promising avenues for T2D treatment and related disorders involving EGFR signaling, bridging systems biology and drug discovery approaches in metabolic disease research.

## Introduction

Type 2 diabetes (T2D) mellitus is a chronic metabolic disorder characterized by insulin resistance and progressive β-cell dysfunction, leading to hyperglycemia[1]. It affects over 462 million people worldwide, with projections suggesting this number could rise to 700 million by 2045[2]. The pathogenesis of T2D involves a complex interplay of genetic, epigenetic, and environmental factors, resulting in a dysregulated glucose homeostasis network[3]. The disease is associated with serious complications, including cardiovascular diseases, neuropathy, nephropathy, and retinopathy, which contribute to increased morbidity and mortality among patients[4].

### Current Understanding and Challenges

Genetic studies, including genome-wide association studies (GWAS), have identified numerous loci associated with T2D susceptibility, shedding light on the molecular mechanisms underlying the disease[5]. These loci implicate various biological pathways, such as insulin signaling, beta-cell function, and lipid metabolism, in T2D pathogenesis[6]. Additionally, the role of adipose tissue, skeletal muscle, and liver in insulin sensitivity and glucose metabolism has been well-established[7]. However, despite these advances, significant gaps remain in our understanding of the intricate molecular mechanisms underlying T2D progression, and the complete regulatory network governing T2D remains in-completely understood.

Current treatments for T2D primarily aim to control blood glucose levels and include lifestyle modifications, oral hypoglycemic agents, and insulin therapy. Commonly used drugs include metformin, sulfonylureas, thiazolidinediones, DPP-4 inhibitors, GLP-1 receptor agonists, and SGLT2 inhibitors[8]. While these treatments can effectively manage hyperglycemia, they often come with limitations such as side effects, patient adherence issues, and a decline in efficacy over time[9]. Furthermore, these treatments do not address the underlying causes of insulin resistance and beta-cell dysfunction, highlighting the need for novel therapeutic strategies.

### Gaps and Opportunities

One of the significant gaps in current T2D research is the incomplete mapping of the gene regulatory networks that drive disease progression. Understanding these networks is crucial for identifying key regulatory nodes and potential drug targets. Recent advances in high-throughput sequencing technologies and bioinformatics have enabled the systematic analysis of gene expression, epigenetic modifications, and protein interactions in T2D (Imamura et al., 2017, Florez, 2017). These approaches can reveal novel regulatory mechanisms and provide insights into the dysregulation of specific genes and pathways in T2D.

Identifying drug targets within these regulatory networks offers the potential to develop therapies that modulate the disease at its root. For example, targeting transcription factors, non-coding RNAs, or signaling molecules that play central roles in T2D-related pathways could lead to more effective and precise treatments[11]. Additionally, repurposing existing drugs based on their effects on these regulatory networks is an emerging strategy that can expedite the development of new treatments[12].

### Objective and Scope

This research paper aims to elucidate the regulatory network involved in T2D and identify potential drug targets and candidates for therapeutic intervention. By integrating data from transcriptomics, proteomics, and functional genomics, we will construct a comprehensive map of the regulatory circuits in T2D. We will also explore the potential of novel and existing compounds to modulate the network, with the goal of uncovering promising candidates for drug development.

In summary, a deeper understanding of the regulatory network in T2D holds promise for identifying new drug targets and developing more effective treatments. Addressing the current challenges in T2D therapy requires a multidisciplinary approach that leverages advances in genomics, bioinformatics, and pharmacology to translate molecular insights into clinical applications.

## Methods

Our study employed a comprehensive approach to investigate the type 2 diabetes (T2D) regulatory network and identify potential drug targets and candidates. The methodology comprised several steps, including differential gene expression analysis, protein-protein interaction network construction, meta-analysis, regulatory network building, functional enrichment analysis, and structure-based virtual screening. This comprehensive approach allowed us to investigate the T2D regulatory network and identify potential drug targets and candidates, providing insights into the molecular mechanisms of the disease and possible therapeutic interventions.

### Differential Gene Expression Analysis

Differential gene expression analyses were performed on the following studies, focusing on gene expression profiles in beta-cell enriched tissue and isolated human islets from T2D and non-diabetic subjects:

- GSE20966[13]: “Gene expression profiles of beta-cell enriched tissue obtained by Laser Capture Microdissection from subjects with type 2 diabetes”
- GSE25724[14]: “Expression data from type 2 diabetic and non-diabetic isolated human islets”
- GSE196797[15]: “microRNA Expression in Normal and Diabetic Pancreatic Islets”

For GSE20966 and GSE25724, the in-house NCBI tool Geo2R was used to perform the analysis, while the web server ExpressAnalyst (https://www.expressanalyst.ca/) was utilized for GSE196797. In studies GSE20966 and GSE196797, raw P-values were selected, whereas adjusted P-values (Benjamini–Hochberg correction) were chosen for GSE25724. A P-value cutoff of 0.05 and positive log fold change values were considered to identify overexpressed genes.

### Protein-Protein Interaction Network Analysis

Overexpressed genes from each study were uploaded separately to the STRING web server (https://string-db.org) to obtain the protein-protein interaction (PPI) networks of the associated proteins. A high confidence coefficient of 0.7 was employed to obtain the networks, which were subsequently analyzed in Cytoscape[16] to generate the top ten hub genes for each study using the CytoHubba[17] plugin, employing both the degree and betweenness metrics.

### Meta-Analysis

A meta-analysis of the three studies was conducted on the ExpressAnalyst server using raw intensity data, relative log expression for normalization, and Combat to adjust for batch effects. The reference group was the control (healthy samples), and the contrast was T2D samples. Raw P-values were selected with a cutoff of 0.05, and Fisher’s method was used for combining P-values in the statistical analysis. Significant genes, including EGFR, which was manually added, were uploaded to STRING to obtain the PPI network with high confidence, and Cytoscape was used to obtain the top 20 hub genes from each metric. The final set of genes was derived from the genes common to both metrics.

### Regulatory Network Construction

The finalist genes were uploaded to the miRWalk[18] online server to check for miRNAs targeting them. The results were filtered by the TargetScan, miRDB, and miRTarBase databases with a minimum score of 0.95, and the three regions (3’UTR, 5’UTR, and CDS) were searched for targets. The genes were also uploaded to the NetworkAnalyst server (https://networkanalyst.ca) to retrieve transcription factors interacting with the genes from the TRRUST, ENCODE, and JASPAR databases. The final regulatory network, comprising genes, TFs, and miRNAs, was constructed using STRING and Cytoscape.

### Functional Enrichment Analysis

Functional enrichment analysis was performed on the genes and TFs using the GeneTrail[19] server. Over-representation analysis was selected for the enrichment algorithm with Benjamini and Yekutieli FDR adjustment and a significance level of 0.04. Minimum and maximum category sizes were set at 2 and 700, respectively.

### Structure-Based Virtual Screening

Given EGFR’s prominence in the regulatory network, we performed a structure-based virtual screening using the PDB structure 4HJO[20], which consists of the crystallized structure of the inactive EGFR tyrosine kinase domain in complex with erlotinib (Figure 1). The structure was retrieved, cleaned of the ligand and non-standard residues, and prepared for docking using UCSF Chimera[21] with default settings for hydrogen addition and Gasteiger charges. Energy minimization was performed with Swiss PDB Viewer[22].

**Figure 1.**
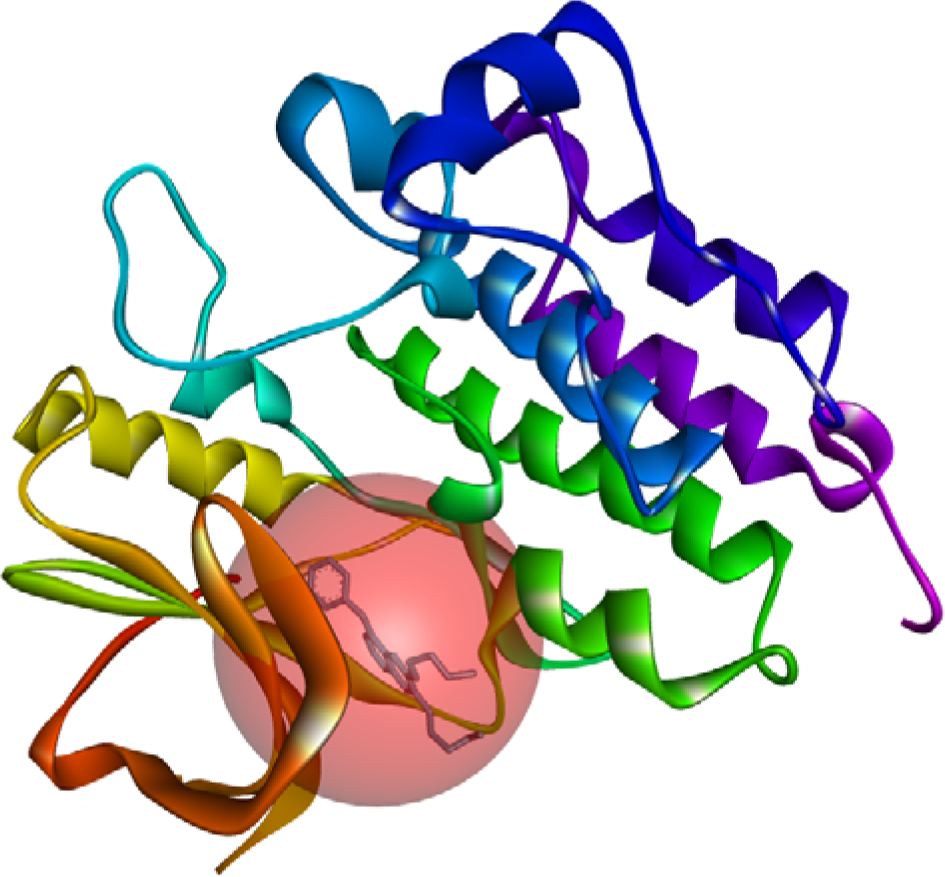
Crystal structure of the inactive EGFR tyrosine kinase domain with erlotinib, obtained through X ray diffraction (PDB Id 4HJO), used for molecular docking. The picture shows the ligand and binding site, and it was obtained from the PDB structure file using Discovery Studio Visualizer.

Ligands were downloaded from ChemDiv[23], Protein Kinases Inhibitors Library (36,324 compounds), and Enamine[24], Kinase Library (64,960 compounds). Prior to docking, ligands were filtered for toxicity properties, drug-likeness, and other criteria using DataWarrior software, and activity prediction using a deep learning model[25] trained on a dataset of tyrosine kinase inhibitors[26] from ChEMBL. Additional drug-likeness filters included Lipinski[27], Ghose[28], Veber[29], Egan[30], and Muegge[31] rules, and structural alerts[32, 33], implemented using Python scripts with the RDKit package.

Filtered compounds were converted to 3D structures, with hydrogens and charges added, and energy minimized using the Amber force field with 5000 steps in Open Babel[34]. Docking was performed with Autodock Vina[35], centered at (28.301, 2.272, 3.051) with a grid size of (20, 30, 20) and an exhaustiveness value of 30, selecting only the highest-scoring poses. Post-docking filters included a binding energy cutoff of -8 kcal/mol, non-inhibition of CYP2C9 and CYP3A4, and BBB permeability. A composite score was calculated using solubility, drug-likeness, drug-score, and binding energy.

### Molecular Dynamics Simulations

The top five scoring compounds from each library were selected for molecular dynamics (MD) simulations of their docked complexes. Simulations were carried out using GROMACS with the CHARMM27 force field and TIP3P solvent, run for 30 ns.

## Results

Our comprehensive analysis of the type 2 diabetes (T2D) regulatory network and potential drug targets yielded several significant findings:

### Regulatory Network Components

We identified ten hub genes that were consistently validated by both degree and betweenness metrics: EGFR, RPS3, AKT1, SKP1, HSP90AB1, BCL2, MED1, SRC, GAPDH, and RPLP0. Additionally, seven transcription factors (TFs) were validated across three databases (TRRUST, ENCODE, and JASPAR): SP1, RELA, PPARG, CREB1, EGR1, JUN, and YY1.

Our miRNA analysis, validated by miRWalk, TargetScan, miRDB, and miRTarBase, revealed five 3’UTR miRNAs (hsa-miR-195-5p, hsa-miR-182-5p, hsa-miR-211-5p, hsa-miR-125b-5p, and hsa-miR-7-5p), one 5’UTR miRNA (hsa-miR-96-5p), and two CDS miRNAs (hsa-miR-181b-5p and hsa-miR-133b). These elements collectively form the T2D regulatory network, as illustrated in Figure 2.

**Figure 2.**
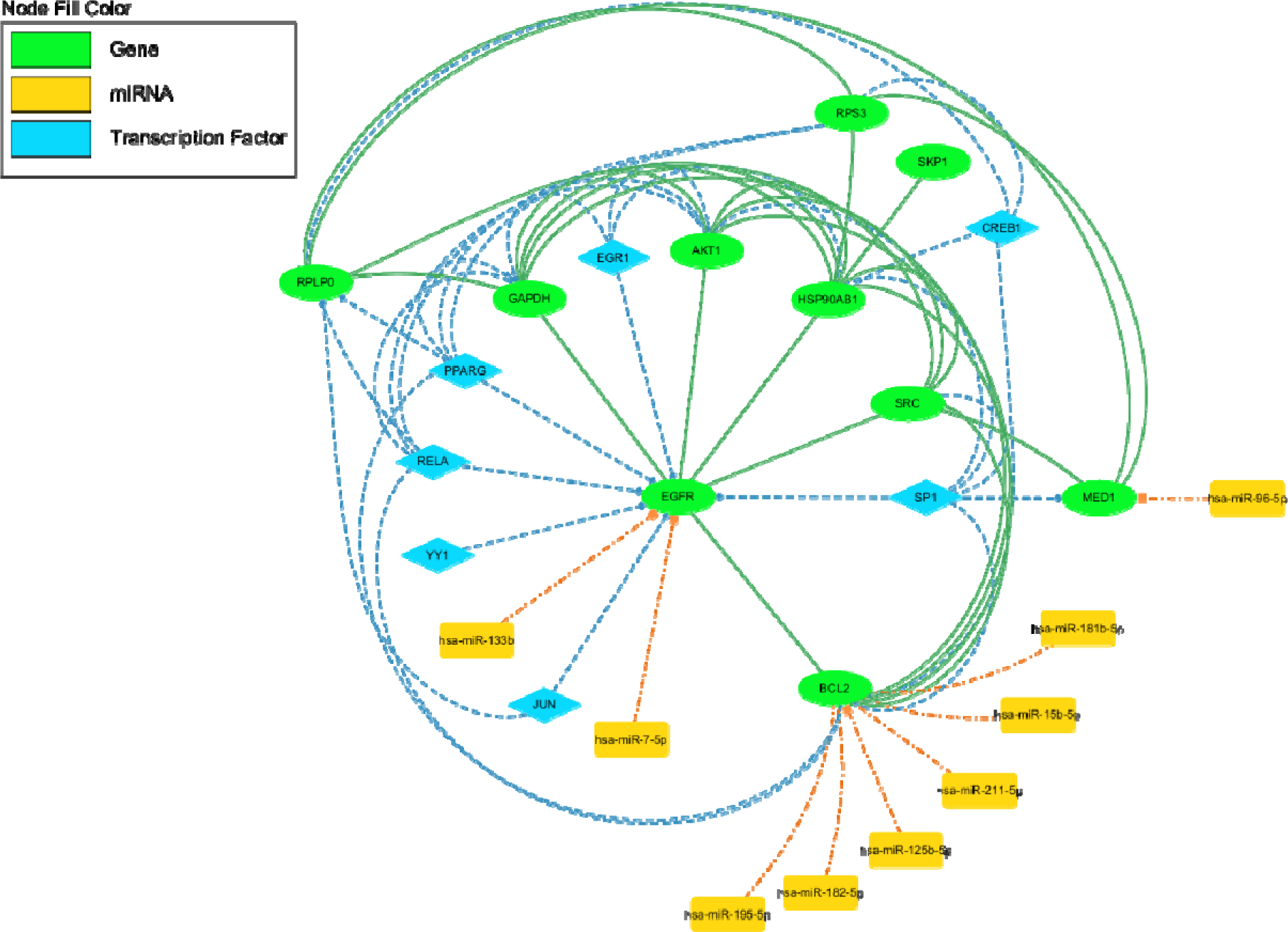
Regulatory Network of Type 2 Diabetes (T2D) involving Genes, Transcription Factors, and miRNAs. The figure depicts the comprehensive regulatory network associated with Type 2 Diabetes (T2D), highlighting the interactions between key genes, transcription factors (TFs), and microRNAs (miRNAs). The network integrates data from differential gene expression analysis, protein-protein interactions, and miRNA-target predictions. Green Nodes represent hub genes identified through degree and betweenness centrality metrics, including EGFR, RPS3, AKT1, SKP1, HSP90AB1, BCL2, MED1, SRC, GAPDH, and RPLP0. Blue diamond nodes represent transcription factors (TFs) validated by TRRUST, ENCODE, and JASPAR databases, including SP1, RELA, PPARG, CREB1, EGR1, JUN, and YY1. Yellow rectangle nodes represent miRNAs targeting the hub genes, validated by miRWalk, TargetScan, miRDB, and miRTarBase databases. These miRNAs include hsa-miR-195-5p, hsa-miR-182-5p, hsa-miR-211-5p, hsa-miR-125b-5p, hsa-miR-7-5p, hsa-miR-96-5p, hsa-miR-181b-5p, and hsa-miR-133b. Solid green edges indicate protein-protein interactions between hub genes. Dashed blue edges represent transcriptional regulation interactions from TFs to target genes. Dotted orange edges represent miRNA-target interactions affecting the expression of hub genes. The network illustrates the central role of EGFR as a primary hub in the T2D regulatory network, interacting with multiple genes, TFs, and miRNAs. The integration of these molecular components provides insights into the regulatory mechanisms underlying T2D and identifies potential therapeutic targets and biomarkers for further investigation.

### Functional Enrichment Analysis

The functional enrichment analysis of the identified genes and TFs highlighted several key biological processes and molecular functions. In the Gene Ontology Biological Process (GO BP) category, the principal enriched processes included:

- Cellular response to organic substance
- Cellular macromolecule biosynthetic process
- Regulation of apoptosis and cell death
- Response to reactive oxygen species (ROS)

For the Gene Ontology Molecular Function (GO MF) category, the main enriched functions were:

- RNA polymerase II DNA-binding transcription factor activity
- Ubiquitin-protein ligase activity
- Enzyme binding
- Organic cyclic compound binding These results are detailed in Figure 3.

**Figure 3.**
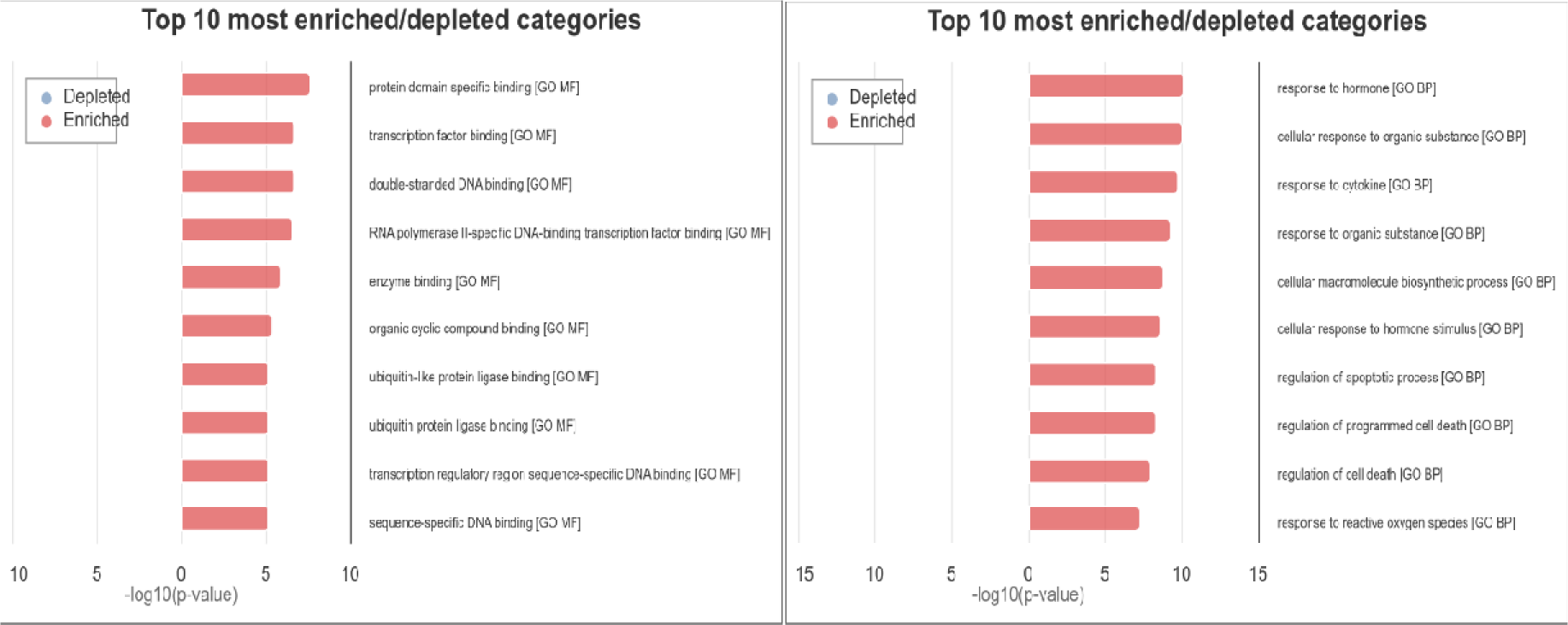
Enriched categories in Gene Ontology (GO) Molecular Function (MF), and Biological Process (BP) for the T2D regulatory network hub genes and TFs, obtained from GeneTrail.

### Identification of EGFR as a Key Therapeutic Target

Differential gene expression analysis consistently identified EGFR as the top hub gene by both degree and betweenness metrics across all three studies analyzed (Figure 4). EGFR also maintained its prominence in the final regulatory network, underscoring its potential as a primary therapeutic target for T2D.

**Figure 4.**
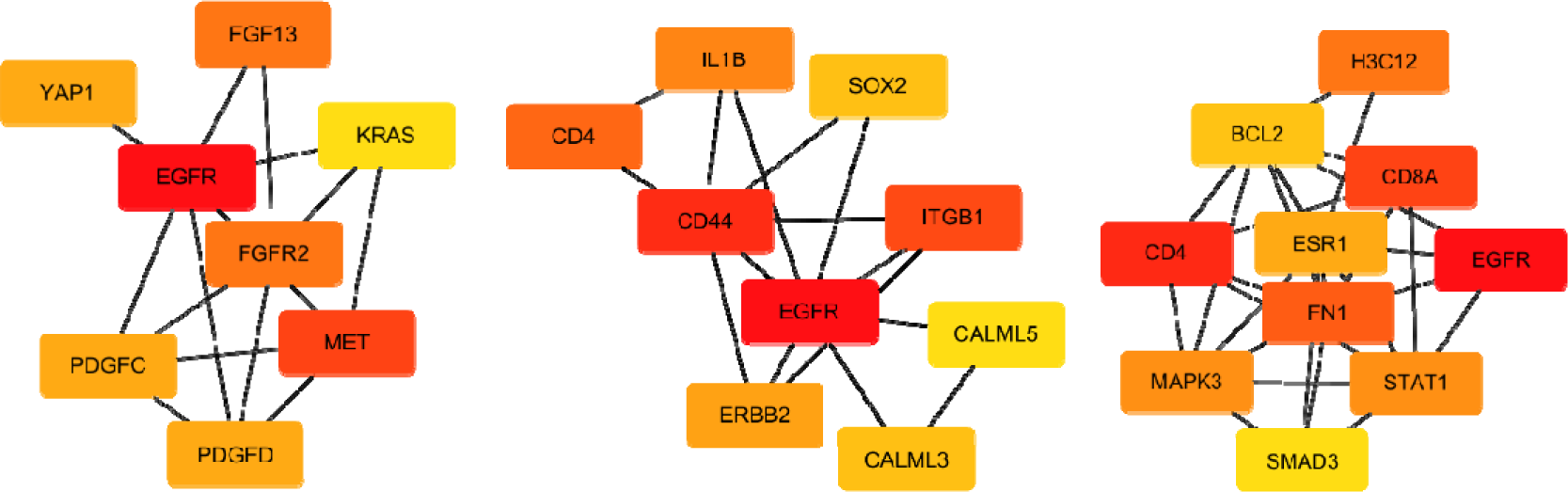
Hub genes with the highest degrees of interaction in the PPI networks of studies GSE196797 (left), GSE20966 (center), and GSE25724 (right). Color code for the degree of interaction: darker red for the highest, lighter yellow for the lowest. The figure highlights the fact that EGFR appears as the top hub gene across the three studies. The result holds if the centrality betweenness metric is used instead.

**Figure 5.**
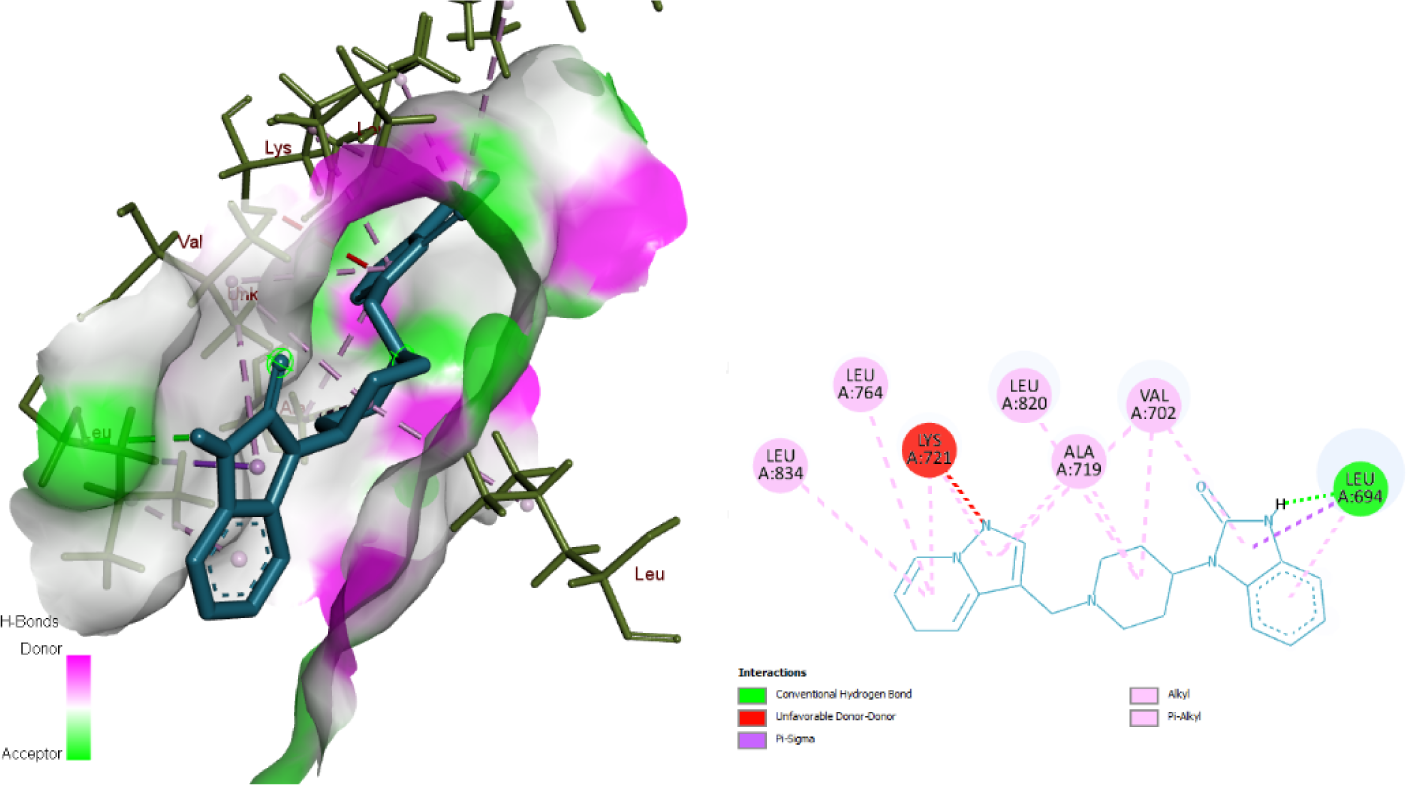
Molecular interactions for the complex EGFR-Z1562794435 (Hit 1). Left: Docked complex three-dimensional diagram, showing the protein’s surface in terms of donor-acceptor hydrogen bonds. Right: Docked complex two-dimensional diagram, showing various interactions between the ligand and the protein.

### Virtual Screening and Molecular Dynamics Simulations

The virtual screening process yielded three candidate compounds with optimal physicochemical and ADMET properties, deemed suitable for further development as lead compounds. These hits included one compound from the Enamine library and two from the ChemDiv library. The docking results for these compounds were validated through molecular dynamics (MD) simulations.

Tables 1-5 provide detailed properties of the hits alongside controls, while Figures 6-11 illustrate the docking poses and MD simulation results. These findings suggest that the identified compounds have promising characteristics for further investigation as potential T2D therapeutics.

**Figure 6.**
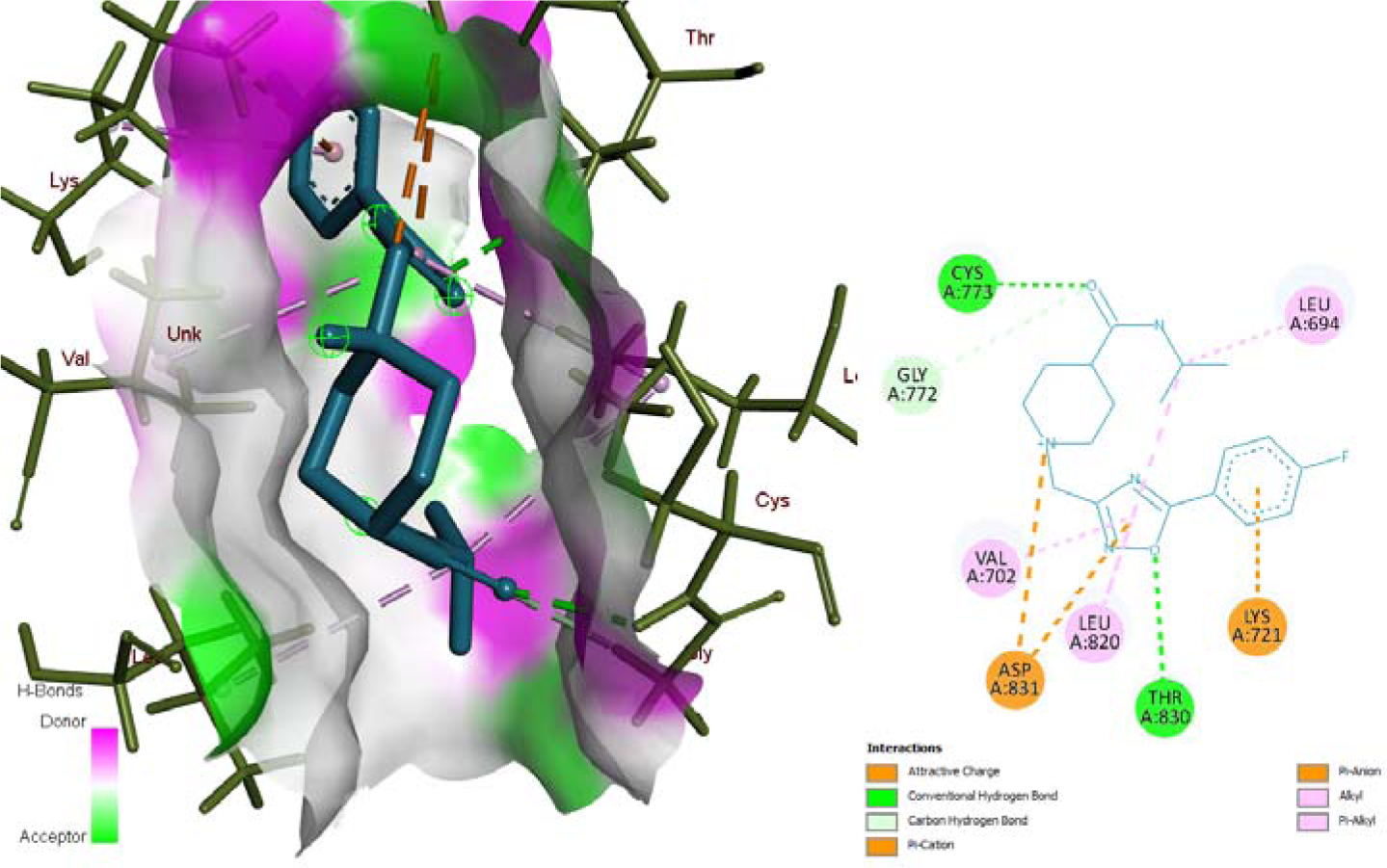
Molecular interactions for the complex EGFR-J065-0160 (Hit 2). Left: Docked complex three-dimensional diagram, showing the protein’s surface in terms of donor-acceptor hydrogen bonds. Right: Docked complex two-dimensional diagram, showing various interactions between the ligand and the protein.

**Figure 7.**
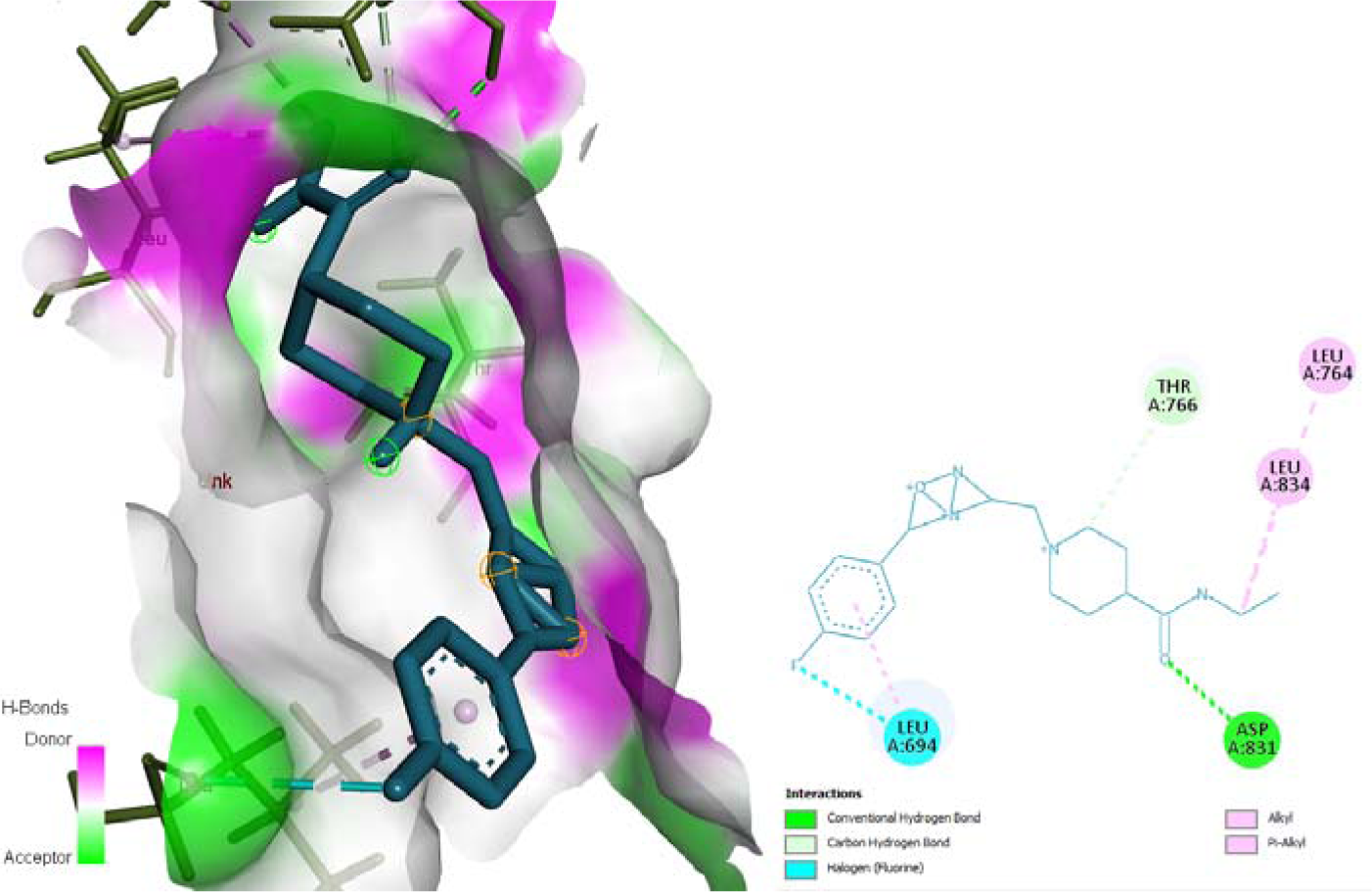
Molecular interactions for the complex EGFR-J065-0145 (Hit 3). Left: Docked complex three-dimensional diagram, showing the protein’s surface in terms of donor-acceptor hydrogen bonds. Right: Docked complex two-dimensional diagram, showing various interactions between the ligand and the protein.

**Figure 8.**
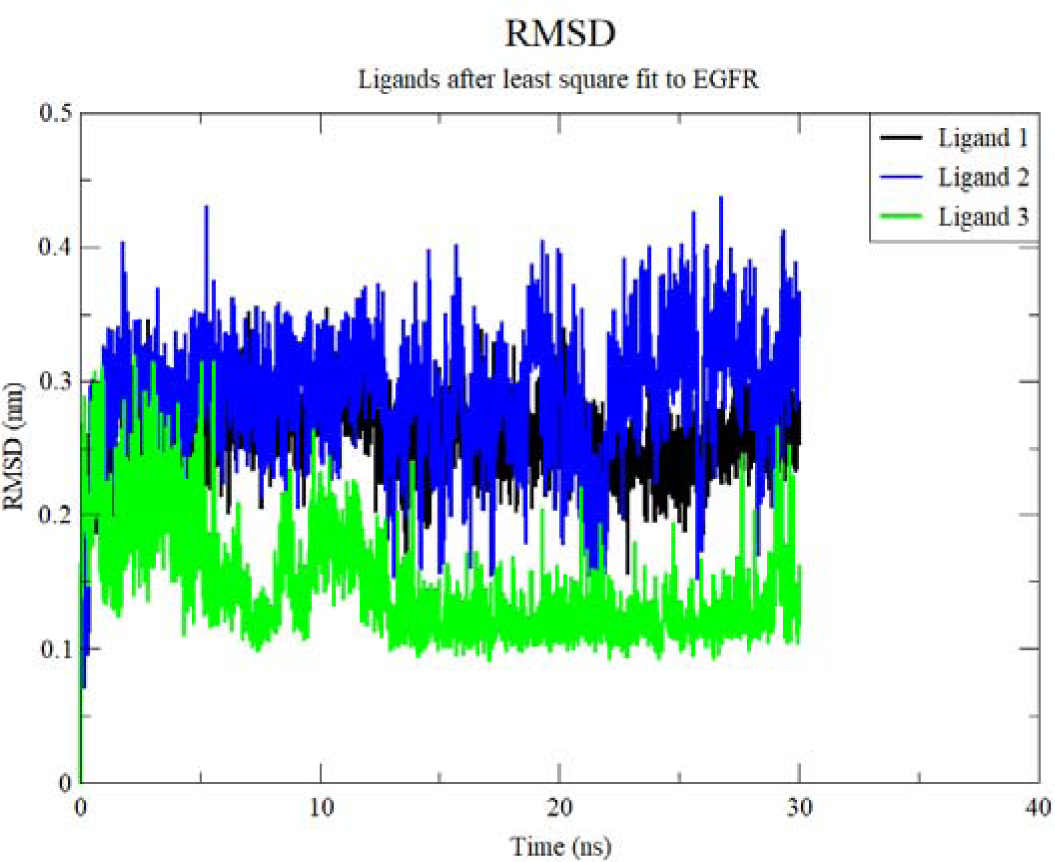
RMSD trajectories of three hit ligands bound to EGFR over a 30 ns molecular dynamics simulation. Ligands’ order is the same as in Table 1. The plot compares the stability of binding poses for Ligand 1 (black), Ligand 2 (blue), and Ligand 3 (green). Ligand 3 demonstrates the most stable binding with the lowest RMSD values, while Ligand 2 shows the highest fluctuations. All ligands maintain RMSD values below 0.5 nm, indicating relatively stable binding throughout the simulation.

**Figure 9.**
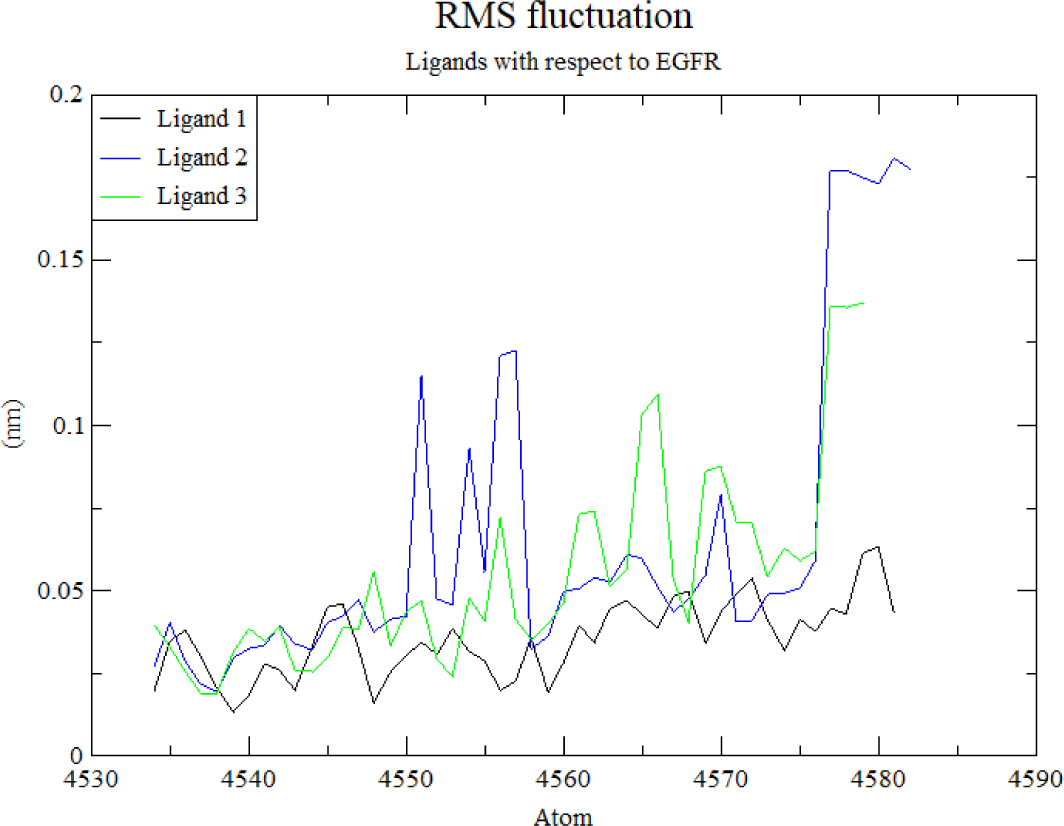
RMS fluctuation profiles of the three hit ligands bound to EGFR. Ligands’ order is the same as in Table 1. The plot compares the atomic flexibility of Ligand 1 (black), Ligand 2 (blue), and Ligand 3 (green) across atoms 4530-4590. Ligand 1 exhibits the most consistent low fluctuations, suggesting stable binding. Ligand 1 and Ligand 3 show higher flexibility in specific regions, particularly around atom 4580, indicating potential areas of increased mobility in their binding poses.

**Figure 10.**
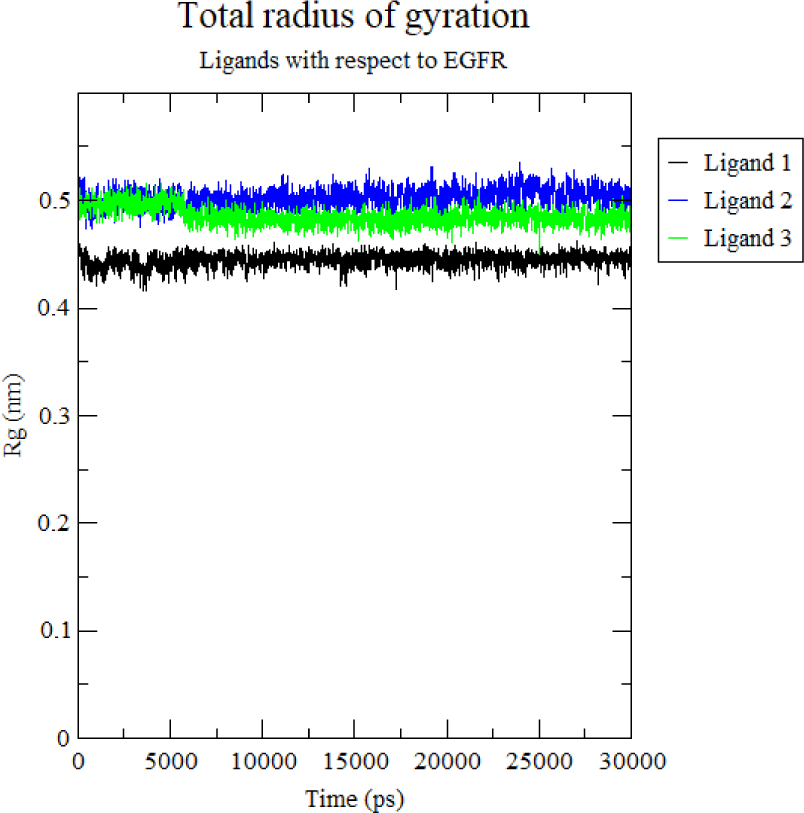
Total radius of gyration for the three hit ligands bound to EGFR over a 30 ns molecular dynamics simulation. Ligands’ order is the same as in Table 1. Ligand 1 (black) exhibits the most compact structure with the smallest radius, while Ligand 3 (blue) shows the largest radius, indicating a more extended conformation. Ligand 2 (green) maintains an intermediate compactness. The stability of these measurements suggests consistent ligand conformations throughout the simulation.

**Figure 11.**
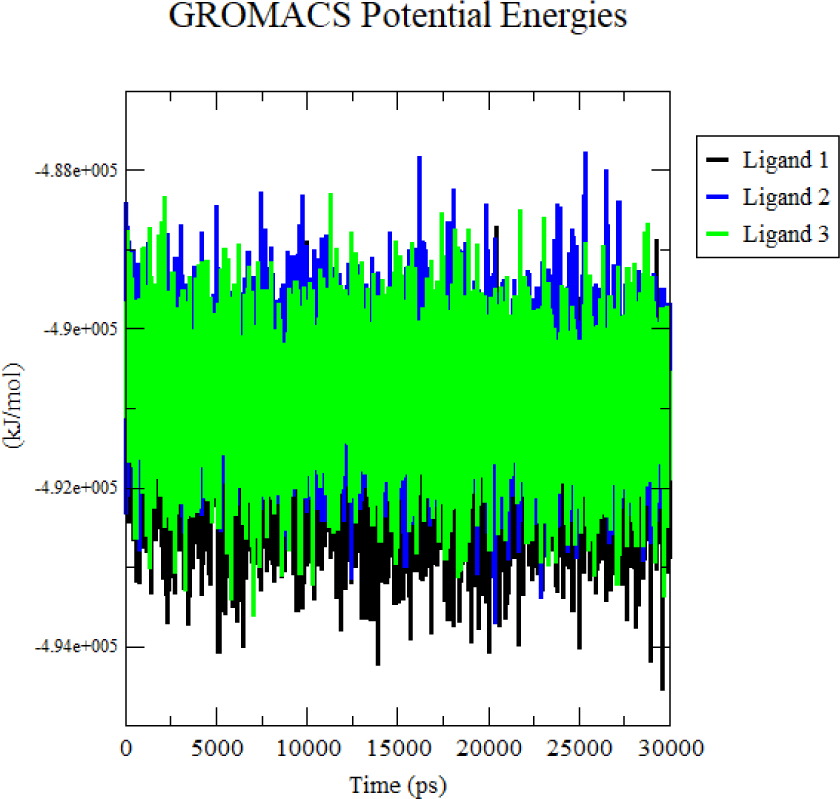
GROMACS Potential Energies for the three hit ligands over a 30 ns molecular dynamics simulation. All ligands exhibit stable energies around -4.9e+005 kJ/mol, with Ligand 1 (black) showing slightly lower energies on average. The consistent energy profiles suggest stable binding conformations for all three ligands throughout the simulation, with minor fluctuations typical of molecular dynamics.

## Discussion

### EGFR as a Key Hub Gene

Despite consistently appearing as the top hub gene in differential gene expression analysis (DGEA) across the three studies, EGFR is only slightly overexpressed with Log FC values of 0.85 in GSE20966, 1.03 in GSE196797, and 0.19 in GSE25724. When false discovery rate (FDR) adjusted P-values are used in the former two studies, very few dysregulated genes result, and the associated PPI network is not significant enough to obtain hub genes. This is why raw P-values were used in those studies.

EGFR’s slight overexpression also explains why it does not show significance in the meta-analysis, as its expression is not strong enough to reach statistical significance in the combined studies. The choice of including EGFR manually in the dysregulated genes resulting from the meta-analysis is justified by its biological significance: the variation in the other hub genes between studies suggests that there might be study-specific factors influencing gene expression, but the repeated identification of EGFR as the top hub gene highlights its importance in the underlying T2D biological processes.

The association between EGFR and T2D has not been conclusively proven, but there is already some evidence for it: EGFR signaling interacts with insulin pathways through shared downstream molecules like PI3K and AKT, crucial for glucose metabolism[36]. Disruption in EGFR signaling can affect insulin action, potentially leading to insulin resistance, a hallmark of T2D.

EGFR’s role in chronic inflammation, a key feature of T2D, further supports its relevance[37]. Some studies have reported elevated EGFR levels in T2D patients, particularly in adipose tissue of obese individuals at high risk for T2D[38]. Moreover, studies have shown that tyrosine kinase inhibitors, particularly ones targeting EGFR such as imatinib, erlotinib and sunitinib, can improve insulin sensitivity and glucose tolerance.[39, 40, 41].

### Functional Enrichment Analysis

The results from functional enrichment analysis confirm the relevance of the obtained regulatory network in T2D biological processes and molecular functions:

- **Apoptosis**: Plays a critical role in the pathophysiology of T2D through its effects on pancreatic β-cells, insulin resistance, and associated complications. Emerging evidence suggests that dysregulation of reactive oxygen species (ROS) metabolism plays a significant role in the pathogenesis and progression of T2D[42, 43]. Excessive ROS production and impaired antioxidant defense mechanisms can lead to oxidative stress, contributing to insulin resistance, beta-cell dysfunction, inflammation, and complications associated with T2D.
- **Cellular Responses to Organic Substances**: There is a significant relationship between cellular responses to organic substances and T2D. This connection primarily involves how cells respond to various organic molecules such as glucose, lipids, and fatty acids, which play critical roles in the development and progression of T2D.
- **Hormonal Regulation**: Hormones play a crucial role in regulating glucose metabolism, insulin sensitivity, and overall energy balance, which are central to the pathophysiology of T2D.
- **Cellular Macromolecule Biosynthetic Process**: Highly relevant to T2D due to its critical role in insulin production, glucose metabolism, lipid synthesis, and overall cellular function. Dysregulation of these processes contributes to the development and progression of T2D.
- **Response to Cytokines**: Relevant to T2D due to their significant roles in inflammation, insulin resistance, adipose tissue function, and pancreatic beta-cell health.
- **RNA Polymerase II Binding**: Plays crucial roles in the transcription of genes involved in insulin production, glucose metabolism, inflammatory responses, and overall metabolic regulation.
- **Ubiquitin Protein Ligase Binding**: Relevant to T2D due to its critical role in regulating insulin signaling, glucose metabolism, beta-cell function, and inflammatory responses. Dysregulation of ubiquitin ligase activity can contribute to the development and progression of T2D.

### Drug Target Considerations

In the context of anti-diabetic drug development, our analysis highlighted the importance of CYP2C9 and CYP3A4 inhibition. These enzymes are crucial for metabolizing key antidiabetic medications like sulfonylureas and meglitinides. To minimize adverse effects and ensure effective drug metabolism, compounds that do not inhibit these enzymes were prioritized in our screening process.

### Promising Drug Candidates

The three hits identified have very good ADMET (absorption, distribution, metabolism, excretion, and toxicity) and drug-likeness properties, outperforming control drugs. Docking results show a significant number of pi interactions contributing to the stability of the complexes, which is further validated by molecular dynamics results. The hits found could also be developed for cancer treatments where EGFR inhibition is important.

### Concluding remarks

While our study provides valuable insights, it’s important to note some limitations. The variation in hub genes across studies suggests the influence of study-specific factors on gene expression. Future research should aim to elucidate these factors and their impact on T2D gene regulation. Additionally, while EGFR’s role in T2D is promising, further experimental validation is needed to conclusively establish its significance as a therapeutic target.

In conclusion, our integrative analysis elucidates key components of the T2D regulatory network, highlight important biological processes and molecular functions involved in the disease, and identify EGFR as a promising therapeutic target. Furthermore, we present three novel compounds that exhibit promising properties and interactions to be developed as potential leads for drug development, both for T2D treatment and other diseases involving EGFR signaling. These findings provide a foundation for future experimental studies and drug development efforts, also opening avenues for exploring multi-purpose therapeutics.

## Authors’ contribution

Ricardo Romero

Conceptualization (Lead), Data curation (Lead), Formal analysis (Lead), Investigation (Lead), Methodology (Lead), Writing – original draft (Lead), Writing – review & editing (Lead).

Abigail Cohen

Data curation (Supporting), Formal analysis (Supporting), Writing – review & editing (Supporting).

The authors declare that they have no known competing financial interests or personal relationships that could have appeared to influence the work reported in this paper.

This research did not receive any specific grant from funding agencies in the public, commercial, or not-for-profit sectors.

This study did not involve any human participants or animal subjects. All data used in this research were obtained from publicly available databases cited in the text, or through computational methods that did not require experimentation on living organisms.

**Table 12.**
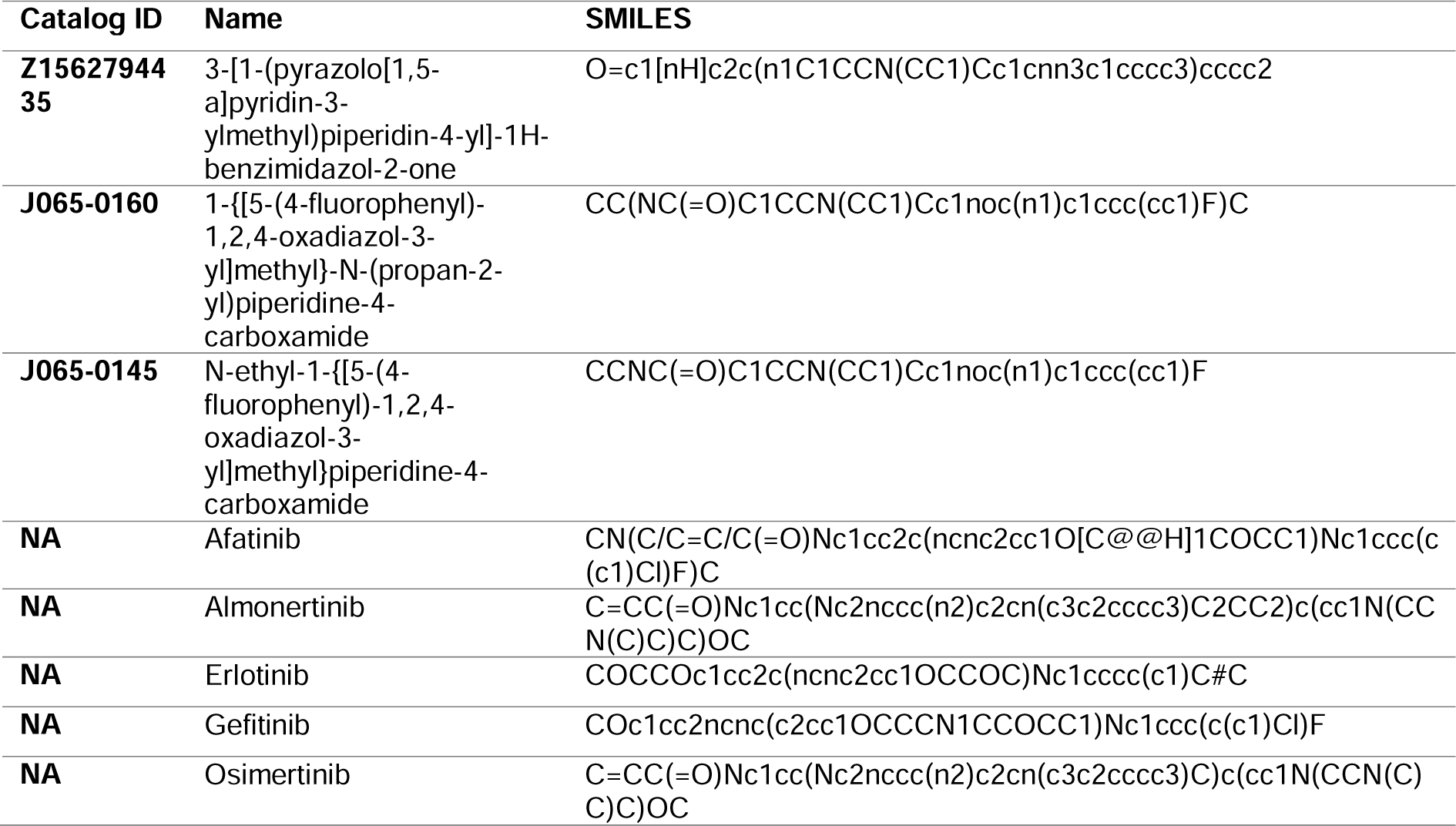
IDs, Names and SMILES of the hits (first entry from Enamine, second and third from ChenDiv) and control drugs (last five entries)

**Table 13.**
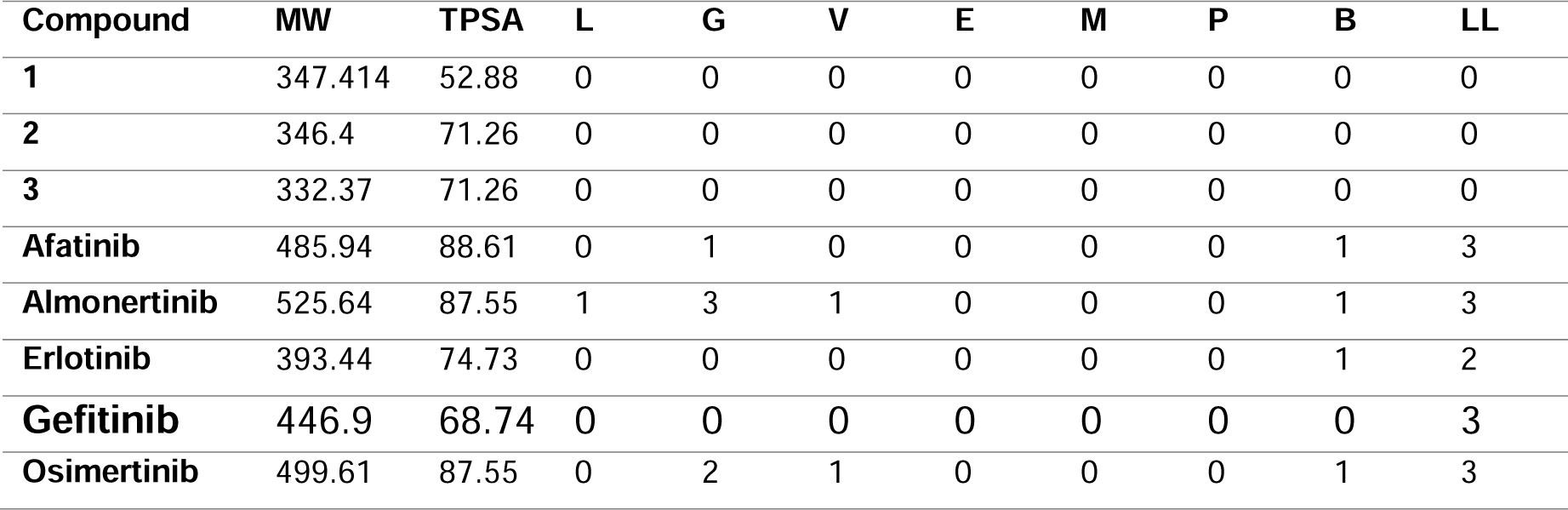
Hits and control druglikeness properties. The numbers 1, 2, 3 refer to the hits in Table 1. Molecular weight (MW) is in g/mol, topological surface area (TPSA) is in LJ^2^. The Lipinski (L), Ghose (G), Veber (V), Egan (E) and Muegge (M) columns indicate the number of violations of the respective rules. The columns P (Pan Assays Interference Structures) and B (Brenk) indicate the number of alerts of the corresponding conditions. The column LL indicates the number of violations of lead likeness properties.

**Table 14.**
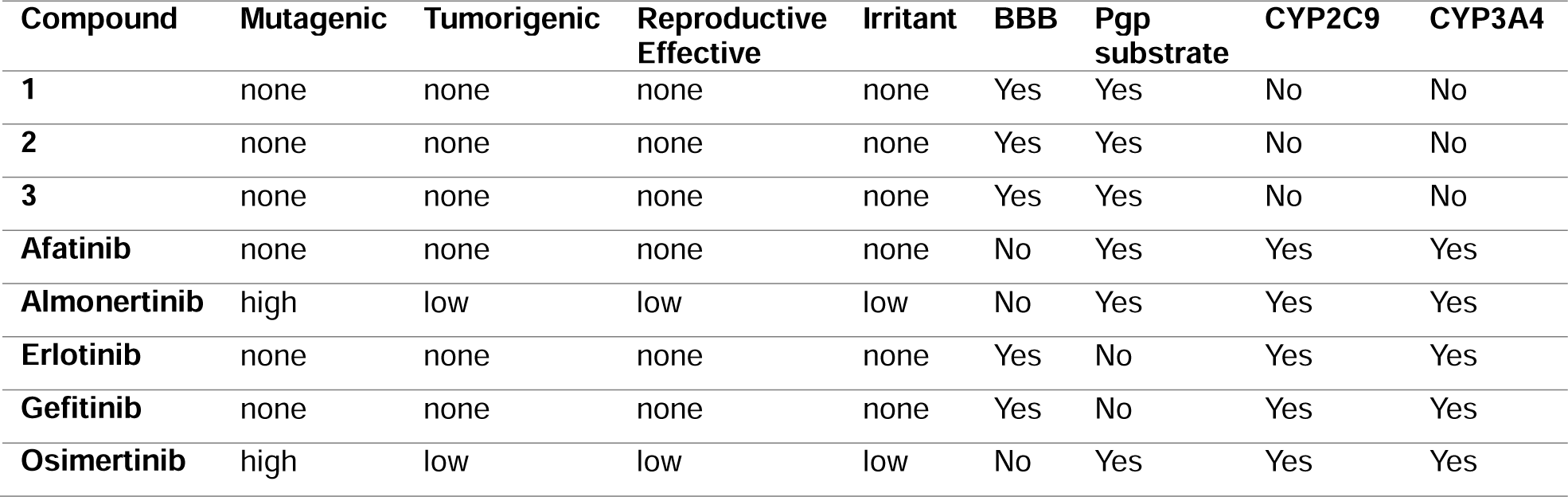
Toxicity and metabolic properties of hits and control drugs. The columns Mutagenic, Tumorigenic, Reproductive effective and Irritant display the corresponding risks according to Data Warrior. BBB indicates brain blood barrier permeability, Pgp substrate indicates transportation by the P-glycoprotein, while the columns CYP2C9 and CYP3A4 indicate whether the compound inhibits the corresponding enzyme from the Cytochrome P450 family. All the compounds show no nasty functions.

| Compound | Mutagenic | Tumorigenic | Reproductive Effective | Irritant | BBB | Pgp substrate | CYP2C9 | CYP3A4 |
| --- | --- | --- | --- | --- | --- | --- | --- | --- |
| 1 | none | none | none | none | Yes | Yes | No | No |
| 2 | none | none | none | none | Yes | Yes | No | No |
| 3 | none | none | none | none | Yes | Yes | No | No |
| Afatinib | none | none | none | none | No | Yes | Yes | Yes |
| Almonertinib | high | low | low | low | No | Yes | Yes | Yes |
| Erlotinib | none | none | none | none | Yes | No | Yes | Yes |
| Gefitinib | none | none | none | none | Yes | No | Yes | Yes |
| Osimertinib | high | low | low | low | No | Yes | Yes | Yes |

**Table 15.** Hits and control drugs scores. The binding energy from docking is given in kcal/mol. Solubility, Druglikeness (OPE), and Drug Score are the scores obtained from OSIRIS Property Explorer. Druglikeness (DW) is the score obtained by Data Warrior. The final score is calculated using the average 1/5(solubility + druglikeness (OPE) + druglikeness (DW) + drug score + |binding energy|). All compounds were classified as active by the deep neural network bioactivity model.

| Compound | Binding Energy | Druglikeness (DW) | Solubility | Druglikeness (OPE) | Drug-score | Final Score |
| --- | --- | --- | --- | --- | --- | --- |
| 1 | -9.4512 | 9.9279 | -3.82 | 10.18 | 0.81 | 5.31 |
| 2 | -8.5006 | 7.5349 | -3.61 | 7.97 | 0.82 | 4.20 |
| 3 | -8.3868 | 7.1996 | -3.23 | 7.64 | 0.86 | 4.20 |
| **Afatinib** | -8.9215 | -0.79682 | -5.48 | -4.11 | 0.24 | -0.25 |
| **Almonertinib** | -8.1943 | -4.9192 | -4.46 | -4.09 | 0.08 | -1.04 |
| **Erlotinib** | -8.0503 | -5.9718 | -3.53 | -6.73 | 0.38 | -1.56 |
| **Gefitinib** | -8.6615 | 0.47937 | -5.06 | -2.62 | 0.28 | 0.35 |
| **Osimertinib** | -8.2775 | -5.4185 | -4.45 | -4.58 | 0.09 | -1.22 |

**Table 16.**
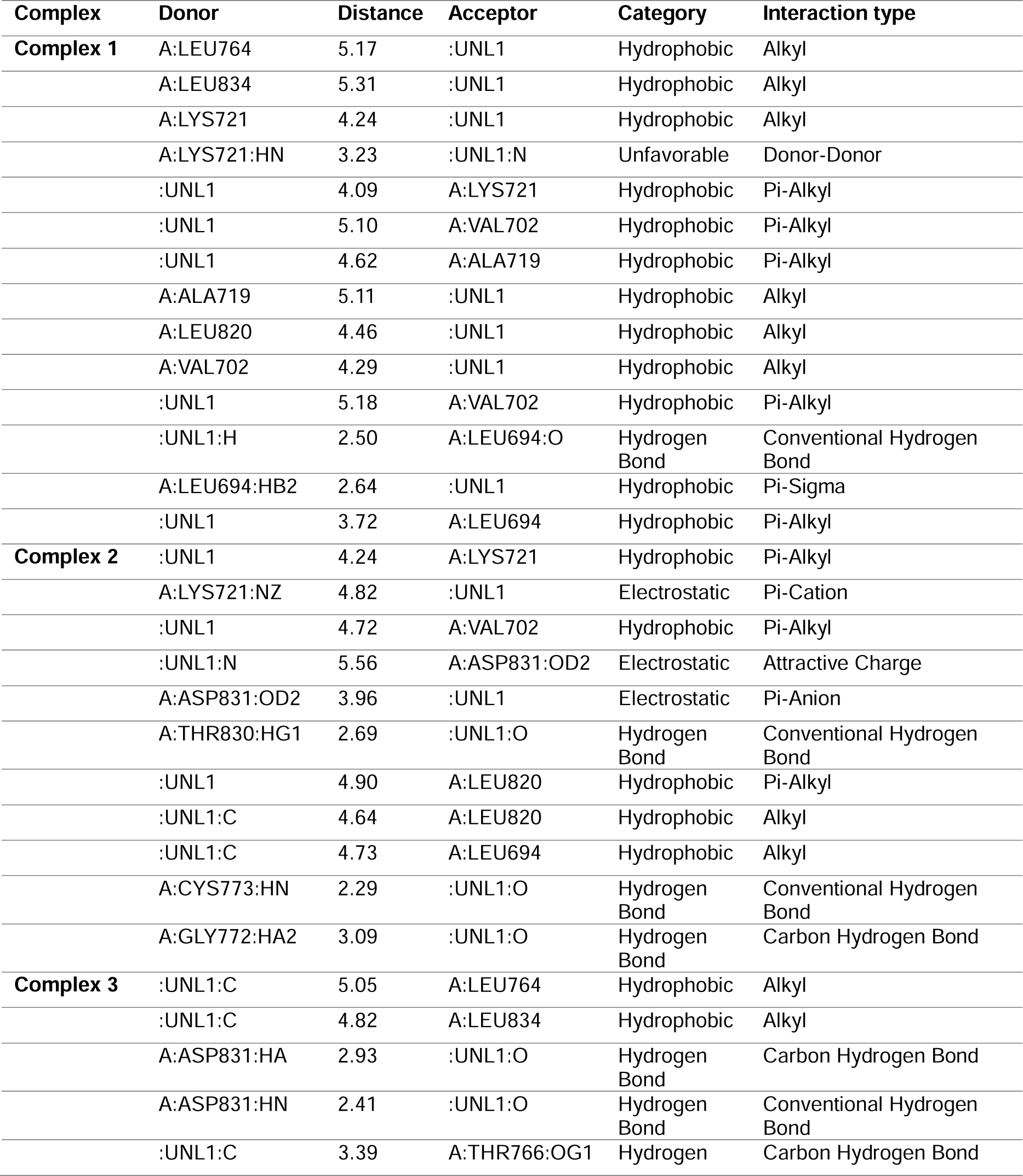
Molecular interactions for the three hits compounds in complex with EGFR. The distance is reported in LJ^2^ and UNL1 refers to the corresponding ligand in each case.

